# Cline coupling and uncoupling in a stickleback hybrid zone

**DOI:** 10.1101/016832

**Authors:** Tim Vines, Anne Dalziel, Arianne Albert, Thor Veen, Patricia Schulte, Dolph Schluter

## Abstract

Strong ecological selection on a genetic locus can maintain allele frequency differences between populations in different environments, even in the face of hybridization. When alleles at divergent loci come into tight linkage disequilibria, selection acts on them as a unit and can significantly reduce gene flow. For populations interbreeding across a hybrid zone, linkage disequilibria between loci can force clines to share the same slopes and centers. However, strong ecological selection can push clines away from the others, reducing linkage disequilibria and weakening the barrier to gene flow. We looked for this ‘cline uncoupling’ effect in a hybrid zone between stream resident and anadromous sticklebacks at two genes known to be under divergent natural selection (*Eda* and *ATP1a1*) and five morphological traits that repeatedly evolve in freshwater stickleback. We used 10 anonymous SNPs to characterize the shape of the zone. We found that the clines at *Eda*, *ATP1a1*, and four morphological traits were concordant and coincident, suggesting that direct selection on each is outweighed by the indirect selection generated by linkage disequilibria. Interestingly, the cline for pectoral fin length was much steeper and displaced 200m downstream, and two anonymous SNPs also had steep clines.

## Introduction

Progress towards speciation can be hindered by hybridization because gene flow erodes allele frequency differences between the populations, and because recombination in hybrids breaks down linkage disequilibria. In general, only alleles under strong selection themselves or close to strongly selected loci are expected to show large allele frequency differences between hybridizing populations (Yeaman & Otto 2011, Yeaman & Whitlock 2011). When strongly selected alleles at different loci come into linkage disequilibrium (i.e. are found together more often than predicted by their respective frequencies), selection tends to act on them as unit, reducing hybrid or immigrant fitness by their combined effects (Bazykin 1969, Barton 1983, Kruuk et al 1999, Barton & de Cara 2009), which helps to maintain divergence between the hybridizing populations. The key processes in achieving further divergence between hybridizing populations are thus a) having increasing numbers of loci with large allele frequency differences between the populations, and b) building stronger linkage disequilibria between the alleles at the diverged loci (Felsenstein 1981, Barton & de Cara 2009, Smadja & Butlin 2011, Flaxman et al. 2013, 2014).

One situation that is particularly informative about the interplay between divergent selection and linkage disequilibria (LD) during the process of speciation is a hybrid zone, where parapatric populations interbreed across a shared boundary (Barton & Hewitt 1985, Harrison 1990). Multiple generations of hybridization between adjacent populations typically produce allele frequency clines at the differentiated loci. The slope and position of each cline is a product of the balance between dispersal and total selection at that locus. Total selection is given by direct selection on the locus plus the sum of indirect selection on diverged loci elsewhere in the genome, weighted by the strength of LD between the focal locus and the other diverged loci (Barton 1983, Barton & Gale 1993). For loci where direct selection is outweighed by indirect selection, all of them will experience approximately the same selection regime, and their clines will cluster together. The clustering of clines in turn increases LD, as ‘pure’ individuals will have alleles characteristic of one or other population at the selected loci, which in turn increases indirect selection (Slatkin 1975, Barton 1983). Clines brought together by strong indirect selection thus tend to be steeper than expected in the zone center (where LD is strongest), and the clines appear to be ‘stepped’ (Szymura & Barton 1991).

By contrast, when direct selection on a locus is stronger than indirect selection, cline shape will reflect the regime imposed by selection on that locus alone. If the direct selection has an ecological component (e.g. the alleles are each favoured on either side of an environmental transition), the cline may be pushed away from the clines at other loci. This scenario is especially likely when the hybrid zone is centered on an ecotone, but the transition points for a subset of environmental variables are displaced away from the boundary between the two environments.

Strong direct selection thus plays two potentially opposing roles: it ensures that allele frequency differences persist despite gene flow (Yeaman & Otto 2011), but it may also push clines apart, reducing linkage disequilibria and thereby weakening the total indirect selection felt by the remaining loci (Nürnberger et al. 1995). Evidence for this ‘cline uncoupling’ effect of strong direct selection is obtained when the clines for genetic loci or phenotypic traits known to be under selection are nearby but do not share cline centers. Conversely, finding that clines for loci under varying types of selection are centered together suggests that direct selection on each locus has been overwhelmed by indirect selection. Further evidence comes from the cline shapes: concordant, stepped clines suggest that indirect selection predominates, while finding a steeper or shallower unstepped cline for a particular locus would indicate that its shape is determined by direct selection alone.

Genetic divergence under gene flow has been well studied, both theoretically (Wright 1931, Bulmer 1972, Yeaman & Otto 2011, Yeaman & Whitlock 2011) and empirically (reviewed in Smadja & Butlin 2011, Feder et al. 2013). By contrast, the circumstances under which clines come together or move apart are less well studied, either theoretically (with the exception of Slatkin 1975, Nürnberger et al. 1995, Bierne et al 2011) or empirically (see Abbott et al. 2013 for a review).

Here, we look for evidence of cline uncoupling in a natural hybrid zone by examining clines for two loci known to be under selection, five ecologically-relevant morphological traits, and 12 anonymous SNPs in a zone between two differentially adapted threespine stickleback populations. In this hybrid zone, one population (‘stream’) inhabits the upper reaches of a small river, Bonsall Creek, throughout the year. The other (‘anadromous’) spends most of the year in the sea and migrates into the lower half of the river to breed over the summer (Hagen 1967). The hybrid zone between the two types is located close to the ecotone between the estuary and the freshwater environment. The estuary itself represents a gradient for many environmental variables, including salinity, tidal range, vegetation, and the community of predators, parasites and competitors. These gradients end abruptly with the transition to freshwater. Sticklebacks can easily swim from one side of the zone to the other, so an active preference for the estuary or freshwater part of the river likely plays a role in structuring the hybrid zone. Moreover, lab crosses between anadromous and stream fish from Bonsall Creek show no evidence of hybrid inviability or infertility (Dalziel et al. 2012, Dalziel & Schulte 2012), and it seems likely that the majority of reproductive isolation in the hybrid zone is extrinsic.

The two selected genes we examine are Ectodysplasin (hereafter *Eda*), and Na^+^, K^+^ ATPase’s catalytic α1 subunit (hereafter *ATP1a1*). *Eda* is located on chromosome IV and is responsible for one of the major phenotypic differences between these populations: stream fish have 4-8 lateral armour plates per side, whereas the anadromous fish have 30-35 (Hagen 1967). The low plated allele at *Eda* has risen to fixation in many freshwater environments (Colosimo et al. 2005), and shows evidence of repeated selection (Mäkinen et al. 2008, Shimada et al. 2011, DeFaveri et al. 2011, Jones et al. 2012, Raeymaekers et al. 2014). *ATP1a1* is located on chromosome I, and is a multi-subunit, membrane bound enzyme that maintains electrochemical gradients by moving K^+^ ions into and Na^+^ ions out of the cell (reviewed by Kaplan 2002). In fish, this enzyme plays a critical role in osmoregulation in both fresh and salt water (reviewed by Evans et al. 2005). Like *Eda,* the *ATP1a1* isoform has an allele that has risen to high frequency in many freshwater stickleback populations (Jones et al. 2006, Shimada et al. 2011) and selection is related to salinity (DeFaveri et al. 2011, 2013; Shimada et al. 2011).

All of the morphological traits we examined have repeatedly evolved after freshwater colonization, and are predicted to be under strong selection. Anadromous populations have long pelvic and dorsal spines that likely defend against gape-limited predators, while freshwater populations typically have shorter spines (Bell et al. 1993, Reimchen 1994, Marchinko 2009). Anadromous fish also have large, long fins that are capable of powering prolonged swimming during migration. Their smaller caudal peduncles are predicted to streamline the fish and reduce drag (Dalziel et al. 2012). Stream fish have repeatedly evolved smaller pectoral fins, which may help fish maneuver in smaller spaces, and deeper caudal peduncles that are predicted to increase burst swimming capacity (Taylor & McPhail 1986). The median fins (which includes the dorsal fin) are involved in maneuvering and force generation during steady swimming (Lauder et al. 2002), and also repeatedly shrink in lakes without predatory fish (Walker 1997; Walker and Bell 2000).

The selection regimes for alleles and traits advantageous in either anadromous or stream fish will not necessarily transition in the same part of the river. For example, selection on loci involved in osmoregulation, such as *ATP1a1*, may favor the stream allele where the stream bed salinity is less than isosmotic (∼13ppt) and the marine allele at higher salinities (Shimada et al. 2011; DeFaveri et al. 2011). By contrast, stream alleles at loci underlying reductions in lateral plates (*Eda*) and other body armor traits (pelvic and dorsal spine length) may only be favored in full freshwater (< 1 ppt), where insects become a significant part of the predator regime. Strong direct selection on these loci and traits would force their clines to be centered in different parts of the river. Our goal in this paper was to thus test whether the clines at body armor loci and traits (*Eda,* pelvic and dorsal spine length), osmoregulation loci (*ATP1a1*) and morphological traits related to swimming capacity (pectoral and dorsal fin length, caudal peduncle depth) have different slopes and centers, which would in turn suggest that strong direct selection has uncoupled the clines at these loci. By contrast, finding that these loci and traits have concordant and coincident clines indicates that indirect selection generated by LD has overwhelmed the presumably strong direct selection and forced the clines into having the same shapes. Either way, our data shed light on the relative roles of direct and indirect selection in promoting (or inhibiting) progress towards speciation in these species.

## Methods

### Data collection

We collected sticklebacks from Bonsall Creek between 15^th^ May and 8^th^ June 2006. The creek is a relatively short (15km) and narrow (∼3m wide) tidal river on the South Eastern end of Vancouver Island, British Columbia (see Hagen 1967). Anadromous stickleback migrate into Bonsall Creek in late spring (Hagen 1967). The majority remains in the tidal part of the creek (<2350m in Figure 1), but a few move further up into the freshwater stretches. Fish that resemble phenotypically pure anadromous sticklebacks (see Figure 3) are rare beyond 2700m upstream. The stream resident population reaches its highest densities upstream of 3300m, but individuals with stream resident phenotypes are common to around 2300m and can occasionally be found as far downstream as 1650m from the sea. Hybrid fish with intermediate phenotypes can be found where the stream resident and anadromous populations overlap, but are most common between 2200m and 2600m.

**Figure 1.**
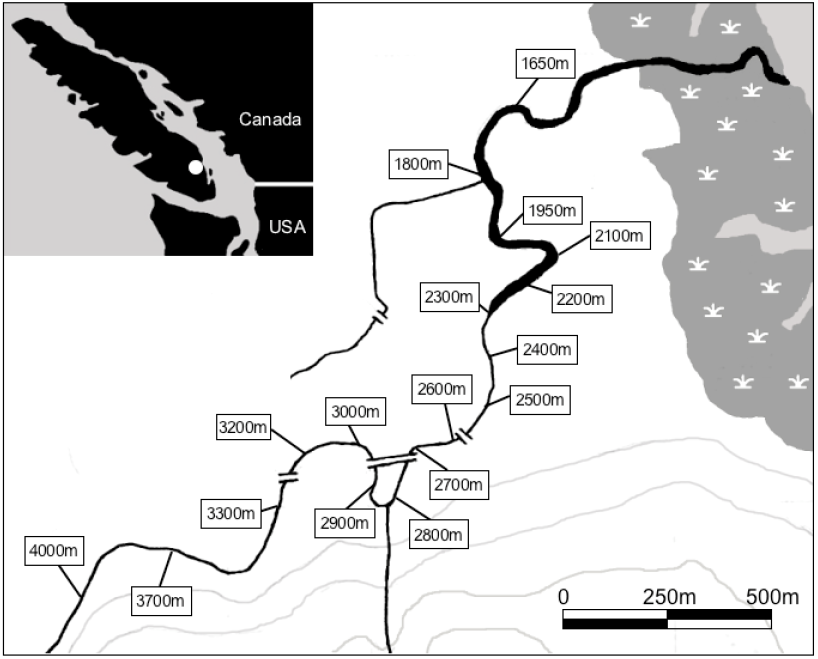
Map of the Bonsall Creek study site on Vancouver Island (white dot on inset) and of the sampling locations along the river itself (main map). The grey shaded area represents salt marsh; the salt finger at high tide reaches 2350m along the creek, although there is some tidal fluctuation in water level at 2400m.

We captured fish at 100 to 150m intervals through the hybrid zone, using 5 minnow traps per site (Figure 1). Fish were killed with an overdose of MS222 anesthetic and frozen on dry ice or preserved in ethanol. We used a YSI 85 Handheld Conductivity and Oxygen meter (YSI Inc., Ohio, USA) to measure the surface and creek bed salinity at each site at low tide and at the peak of the highest tide during sampling period (11pm on 14^th^ June 2006), and found the limit of the salt finger to occur 2350m from the sea (at 48°52'48.46"N, 123°40'26.93"W).

### Morphological data

We selected ten morphological traits that have been found to repeatedly diverge between anadromous and freshwater sticklebacks (Hagen & Gilbertson 1972). Ten marine and ten stream individuals from each side of the hybrid zone were measured for standard length, body depth, head depth, eye diameter, dorsal fin length, caudal peduncle depth, pectoral fin length, caudal peduncle width, left pelvic spine length, and second dorsal spine length. Eye diameter, head depth and body depth showed no consistent differences between marine and stream fish, even when each trait was regressed onto standard length to remove the effects of size, and these traits were discarded. The remaining seven traits were measured on the 428 ethanol preserved fish. We measured standard length and dorsal and pelvic fin length with calipers, the length of the second dorsal and pelvic spine and the caudal peduncle depth and width with an ocular micrometer fitted to a binocular microscope. As standard length showed a strong relationship with all traits, we took the residuals from a regression of each trait on standard length. These residuals were then log transformed to improve their normality.

### Genetic data

We extracted DNA from tail fin tissue from all 428 fish using the protocol in Peichel *et al.* (2001); the DNA was resuspended in 50μl of double distilled water and stored at - 20°C.

#### Ectodysplasin (Eda)

The *Eda* locus is located on linkage group IV. We chose a T/C SNP at position 421 of an amplicon spanning the 7^th^ and 8^th^ exon of the *Eda* gene (Colosimo et al. 2005) for genotyping. The primers are given in Table S1.

#### Na^+^,K^+^ ATPase subunit α1a (ATP1a1)

There are two Na^+^, K^+^ ATPase α1a paralogs in the stickleback genome, which are found in tandem within approximately 9,000 bp of each other on contigs 7064/7065 and 7066 within Linkage group/Chromosome 1. We studied the isoform on contig 7066, which is the same isoform studied by Jones et al. (2006) and Barrett et al. (2008). Our initial genotyping of the anadromous and stream alleles at the 16^th^ to 18^th^ intron of this gene followed the procedure in Jones et al. (2006), but as these sites are not variable in our populations we could not achieve allele-specific amplification. We therefore sequenced this genomic region in 10 pure stream (above 4000m) and 10 pure anadromous fish (from 1650m). We found a number of other diagnostic SNPs within this region, and selected an A/G SNP at position 446 of the amplicon (see Table S1 for sequences and primers).

#### Anonymous SNPs

We began with the list of 25 loci used to construct the phylogenetic tree in Colosimo *et al.* (Figure 3C in Colosimo et al. 2005). We excluded loci on the same linkage group as Eda (LG 4) or Na^+^, K^+^ ATPase (LG 1), although loci with an unknown location were retained. We then tested the primers pairs from Colosimo et al. (2005) on ten stream (4000m) and ten anadromous (1650m) fish from Bonsall Creek. Of the 16 loci that amplified successfully in both populations, we retained 13 loci that had a SNP minor allele frequency greater than 0.1 in at least one of the samples.

These sequences of these 13 loci (Supplementary Table 1) were used by the McGill University and Génome Québec Innovation center to design a custom genotyping assay using Sequenom® iPLEX®Gold Genotyping Technology. Genomic DNA from all 428 fish was quantified and diluted to 20 ng/μl and at least 30μl of each sample were sent on dry ice to the McGill University and Génome Québec Innovation center for genotyping.

**Table 1.** Sample sizes, mean allele frequencies for *Eda* and *ATP1a1*, mean and variance for the Hybrid Index (HI) as calculated from *Eda*, *ATP1a1*, P3D05, and P6A10. Also shown are mean body length and the mean logged residuals from the morphological traits, by site. The limit of the salt finger for the highest tide during the study period was 2350m from the sea.

| Distance from sea (m) | N | *Eda* anadromous allele freq | *ATP1a1* anadromous allele freq | Mean Hybrid Index | Var in Hybrid Index | Body length (mm) | Dorsal fin | Peduncle depth | Pectoral fin | L Pelvic spine | 2<sup>nd</sup> dorsal spine |
| --- | --- | --- | --- | --- | --- | --- | --- | --- | --- | --- | --- |
| 1650 | 28 | 0.80 | 0.83 | 0.81 | 0.04 | 51.0 | 0.58 | -0.10 | 0.31 | 0.79 | 0.60 |
| 1800 | 22 | 0.81 | 0.76 | 0.81 | 0.04 | 48.3 | 0.26 | 0.07 | 0.79 | 0.54 | 0.27 |
| 1950 | 20 | 0.75 | 0.68 | 0.74 | 0.03 | 47.7 | 0.59 | -0.07 | 0.55 | 0.76 | 0.37 |
| 2100 | 43 | 0.71 | 0.69 | 0.71 | 0.06 | 46.9 | 0.33 | -0.06 | 0.20 | 0.45 | 0.25 |
| 2200 | 41 | 0.56 | 0.48 | 0.60 | 0.07 | 47.1 | 0.18 | -0.02 | -0.17 | 0.25 | 0.22 |
| 2300 | 35 | 0.57 | 0.45 | 0.53 | 0.11 | 45.4 | 0.14 | 0.00 | 0.00 | 0.25 | 0.12 |
| 2400 | 24 | 0.37 | 0.48 | 0.50 | 0.06 | 46.4 | 0.11 | 0.00 | -0.36 | 0.11 | 0.03 |
| 2500 | 23 | 0.42 | 0.37 | 0.40 | 0.07 | 44.1 | -0.12 | -0.01 | 0.08 | -0.14 | 0.02 |
| 2600 | 31 | 0.23 | 0.28 | 0.28 | 0.06 | 46.9 | -0.20 | 0.05 | -0.02 | -0.12 | -0.07 |
| 2700 | 35 | 0.24 | 0.26 | 0.24 | 0.06 | 47.5 | -0.17 | 0.04 | 0.00 | -0.39 | -0.27 |
| 2800 | 11 | 0.38 | 0.15 | 0.23 | 0.04 | 43.1 | -0.29 | 0.04 | 0.11 | -0.40 | -0.34 |
| 2900 | 27 | 0.19 | 0.15 | 0.14 | 0.03 | 45.1 | -0.34 | 0.06 | -0.01 | -0.36 | -0.26 |
| 3000 | 25 | 0.06 | 0.08 | 0.08 | 0.01 | 49.1 | -0.45 | 0.10 | -0.10 | -0.81 | -0.46 |
| 3200 | 9 | 0.00 | 0.17 | 0.04 | 0.00 | 47.3 | -0.60 | 0.02 | -0.25 | -0.98 | -0.53 |
| 3300 | 17 | 0.00 | 0.09 | 0.08 | 0.02 | 46.0 | -0.42 | 0.02 | -0.19 | -0.71 | -0.29 |
| 4000 | 21 | 0.00 | 0.00 | 0.02 | 0.00 | 46.6 | -0.49 | 0.00 | -1.10 | -0.62 | -0.42 |

### Statistical methods

#### Cline fitting: genetic loci

The clines at *Eda* and *ATP1a1* and the anonymous SNP loci were characterized with the program CFit version 7 (Gay et al. 2008). This program uses a Metropolis algorithm to find the cline shape that best fits the available data. The program fits two basic parameters that describe a sigmoid cline, cline slope *s* and cline center *c*, and also allows for deviations from Hardy-Weinberg equilibrium (fitted as F_IS_) within each site. Initial tests found no support for including F_IS_ at any of the genetic loci and we do not consider it further.

We also fitted a more complex stepped cline model, where the center part of the cline remains sigmoid, but exponential curves are permitted for each of the tails. This model has six parameters: slope *s*, center *c*, two locations where the exponential tails begin (*xpos1*, *xpos2*) and the two slopes of the exponential curves (*tslope1* and *tslope2*); the latter two range between 0 and 1 and are the proportion by which the slope of the sigmoid curve is reduced. Note that *xpos2* and *tslope2* are the parameters for the anadromous side of the hybrid zone.

#### Cline fitting: morphological data

CFit requires morphological traits to be at their maximum at the left hand side (i.e. anadromous) side of the hybrid zone. This is true for the logged residuals of dorsal fin length, left pectoral fin length, pelvic spine length, and second dorsal spine length. However, the logged residual of caudal peduncle depth is greatest in stream fish, and so we used the negative of the logged residuals in the CFit analysis for this trait.

We used a simple clinal model to characterize the morphological data across the hybrid zone (Gay et al. 2008). Observed measurements were assumed to be drawn from a normal distribution N(*μ*_*x*_, *σ*_*x*_), where *μ*_*x*_ and *σ*_*x*_ are functions of distance *x* across the hybrid zone. More specifically, *μ*_*x*_ was modeled as *μ*_min_+(*μ*_max_−*μ*_min_) *p*_*x*_, where *μ*_min_ and *μ*_max_ measure the mean in the stream and anadromous population, respectively, and *p*_*x*_ is the sigmoid function of distance (with maximum slope *s* and center *c*). The variance *σ*_*x*_ was modeled as 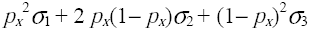 to account for differences among individuals from either the anadromous or the stream populations (*σ*_1_ and *σ*_3_, respectively) and between either pure population and hybrids from the zone center (*σ*_2_). This model assumes that in each location, the distribution of each trait is unimodal (drawn from a simple normal distribution). As with the genetic data, parameters were estimated by maximum likelihood using CFit 7.

For each trait, we compared the fit when the trait variance is constant through the zone (*σ*_1_=*σ*_2_=*σ*_3_) to a model where *σ*_1_, *σ*_2_ and *σ*_3_ were allowed to vary independently. The best fit from this comparison (as determined by comparing the Akaike Information Criteria, Akaike 1973) was then retained for the cline comparison step described below. As with the genetic data, we fitted the stepped cline to the morphological data as well, leading to a model with a maximum of eleven independent parameters.

#### Comparing cline shapes

We tested for coincidence and concordance between clines by forcing them to have the same center and/or slope, respectively, and comparing the AIC values of these models to models where each cline was fitted independently. We considered a difference of 2 AIC units between models an indication of a ‘significant’ difference, with the caveat that a difference of 7 to 10 AIC units is needed before concluding that the worse model has essentially no support (Burnham and Anderson 2002).

#### Cline Analysis with the Hybrid Index

When genetic loci are more or less fixed on either side of the zone they can be treated as diagnostic, and can thus used to calculate a Hybrid Index (HI) for each individual. The Hybrid Index is calculated by counting the number of anadromous alleles across the four loci with large differences in allele frequency between the two populations (*Eda*, *ATP1a1*, P3D05, and P6A10, see results). We can use the variance in HI in the center of the zone to estimate linkage disequilibria 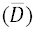 and hence the dispersal rate *σ* (p23 of Barton & Gale 1993). Since cline width is a product of the relationship between dispersal and selection, we can use this information to estimate the strength of selection acting on the genetic loci and the morphological traits.

## Results

### Sample sizes

We genotyped 428 adult (>40mm) fish from 16 sites through the hybrid zone at *Eda*, *ATP1a1*, and 13 anonymous SNP loci. These data are available on Dryad (doi:XXXX). Locus P7A09 failed to amplify in most anadromous individuals and was excluded from the subsequent analyses. Amplification success was otherwise very good, with an average of 9.7 (2.2%) missing genotypes per locus. Two of the anonymous loci (P3A06 and P6B12) showed no allele frequency differences across the hybrid zone, and were also excluded at this stage.

We also obtained morphological data for these 428 fish (Table 1 and Dryad doi:XXXX). We discovered that variation in caudal peduncle width was found to result from two confounding sources: anadromous fish had a bony ‘keel’ which adds 1-2mm to the total width of the peduncle, but this feature is not present in stream individuals. However, when the keel is ignored, stream fish had a wider caudal peduncle than anadromous fish. As the total width of the peduncle could not be expected to fit a simple cline model it was discarded at this stage.

### Comparing cline centers and slopes

Our central question is whether the clines for *Eda*, *ATP1a1*, and the five morphological traits have the same center and slope. We therefore fit a model where these seven clines were constrained to share the same slope and center, and compared it to one where slope and center were allowed to vary independently. A joint center (2298m) and slope (-2.83) was a better fit (AIC = 4206.9 for joint clines vs 4212.8 for independent clines). The parameters for the individually fitted clines at all genetic loci are given in Table 2a.

**Table 2a.**
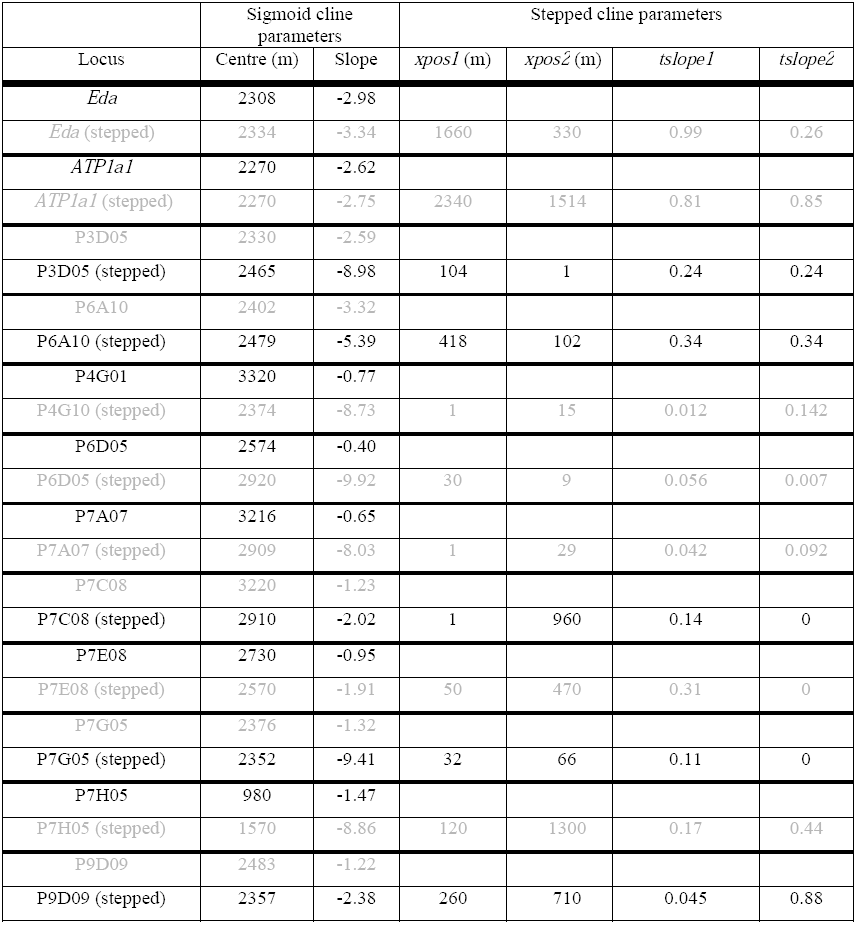
Parameters for the individually fitted two parameter and stepped clines for all genetic loci. Note that the *xpos1* and *xpos2* variables give the number of meters upstream and downstream (respectively) from the center where the exponential tails begin. The best fit cline is highlighted in bold

The cline for pectoral fin length is quite different from the other morphological traits (see above, Figure 3c and S1), and correspondingly the model fit is somewhat improved when it is allowed to have an independent slope and center (AIC = 4201.4 when pectoral fin is independent vs AIC = 4206.9 when it is not). The joint cline for these six traits and loci was at 2296m and had a slope of -2.87.

The clines for six of the seven selected traits and loci are thus centered together in the same part of the river, about 50m downstream from the upper limit of the salt finger at 2350m (Figure 4). We next tested whether these six clines share the same slope. We fit a model where these six clines are constrained to share the same center, but slope is allowed to vary independently. For comparison with the models above, the cline for pectoral fin length was included but had its own slope and center. This model was not an improvement over the model where all six clines shared the same center and slope (AIC = 4201.4 for joint slopes vs 4207.6 for independent slopes). As mentioned in the introduction, it is possible indirect selection is responsible for the shape of most clines, but individual traits or loci may be under strong direct selection and therefore much steeper. We therefore fit a series of model where five of the clines had the same slope and one cline had an independent slope, with all clines sharing the same center. None of these models was a significant improvement over the simpler model where all six clines are constrained to share the same slope and center.

**Figure 4.**
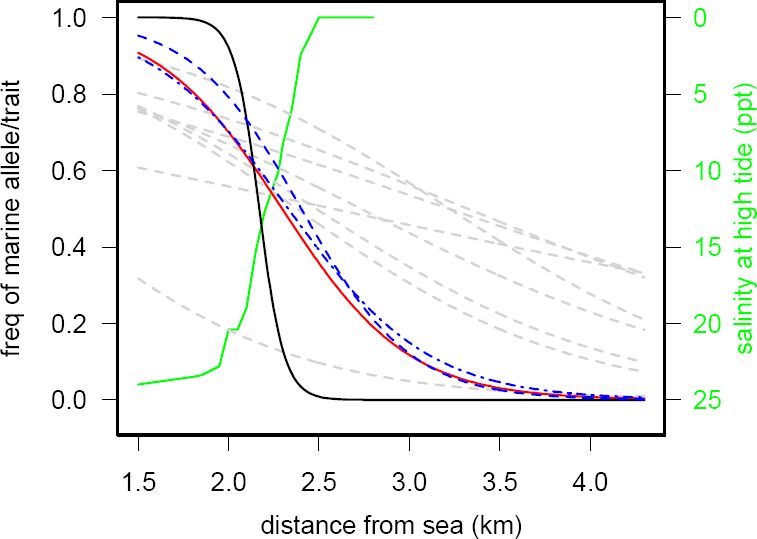
The best-fit clines in the hybrid zone. Salinity at high tide (green line) is plotted on an e scale on the right hand axis. The black line is the cline for pectoral fin length. The red the joint cline for *Eda*, *ATP1a1*, dorsal fin length, caudal peduncle depth, pelvic spine length, and 2^nd^ dorsal spine length. The sigmoid cline for P3D05 is shown with a dot-dash blue line and the sigmoid cline for P6A10 with a dashed blue line. The grey dotted lines are the best-fit sigmoid clines for the remaining eight anonymous SNPs.

The clines for two of the anonymous SNPs (P3D05 and P6A10) are also steep and centered near the joint cline for the selected traits and loci (Table 2a, Figure 4). As a *post hoc* test we fit models where each of these was forced to share the same slope and center as the joint cline. We found some evidence that P3D05 could be included with the joint cline (AIC = 4090.1 when P3D05 is included vs AIC = 4091.5 when P3D05 has its own slope and center). However, the model fit was worse when P6A10 was included with the joint cline (AIC = 4074.26 when P6A10 is included vs AIC = 4062.3 when P6A10 has independent slope and center). This is largely because the cline at P6A10 is located 100m upstream of the joint cline (at 2401m): forcing P6A10 to have the same center as the joint cline is a significantly worse model fit than forcing it to have the same slope (AIC = 4071.3 when P6A10 has joint center but independent slope vs AIC = 4062.8 when it has joint slope but independent center).

### Fitting stepped clines

There was a worsening in model fit when we fitted a stepped cline for both *Eda* and *ATP1a1* relative to the fit of the basic sigmoid cline (*Eda*: AIC=681.6 for sigmoid vs 685.5 for stepped; *ATP1a1*: 713.6 for sigmoid vs 721.36 for stepped). Similarly, a stepped cline model was a significantly worse fit for dorsal fin, peduncle depth, left pelvic spine or 2^nd^ dorsal spine. Pectoral fin length had a much steeper slope (-14.5) and a cline center at 2170m, about 130m downstream of the other traits (Table 2b and Figure 3c). The distribution of pectoral fin phenotypes in the hybrid zone was also quite different from the other four traits. First, fish from the furthest point upstream had proportionally much smaller fins than any of the other sites in the hybrid zone. Second, fish with proportionally very long pectoral fins (as is typical of anadromous fish) were very rare upstream of 2100m (see Figure 3c). We therefore used CFit7 to assess the fit of more complex cline models (see Gay et al. 2008 and the CFit7 manual for details). The best-fit model required 18 parameters and is shown in Figure S1.

**Table 2b.**
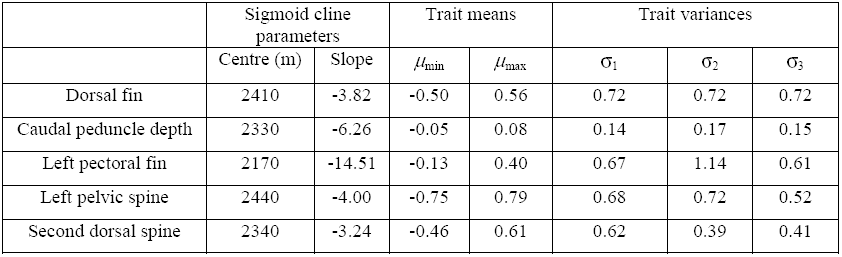
Parameters of the individually fitted clines for all morphological traits. As there was no support for stepped clines for any morphological trait, only the variables for sigmoid clines are shown.

|  | Sigmoid cline parameters |  | Trait means |  | Trait variances |  |  |
| --- | --- | --- | --- | --- | --- | --- | --- |
| | Centre (m) | Slope | $\mu_{\min}$ | $\mu_{\max}$ | $\sigma_1$ | $\sigma_2$ | $\sigma_3$ |
| Dorsal fin | 2410 | -3.82 | -0.50 | 0.56 | 0.72 | 0.72 | 0.72 |
| Caudal peduncle depth | 2330 | -6.26 | -0.05 | 0.08 | 0.14 | 0.17 | 0.15 |
| Left pectoral fin | 2170 | -14.51 | -0.13 | 0.40 | 0.67 | 1.14 | 0.61 |
| Left pelvic spine | 2440 | -4.00 | -0.75 | 0.79 | 0.68 | 0.72 | 0.52 |
| Second dorsal spine | 2340 | -3.24 | -0.46 | 0.61 | 0.62 | 0.39 | 0.41 |

Fitting a stepped cline led to an improvement in model fit for three of five anonymous SNP loci but the fit was hardly improved for the other two. A stepped cline with *tslope1* = *tslope2* was a better fit for P6A10 (AIC = 674.4 for stepped vs 683.2 for sigmoid). However the improvement in fit of a stepped cline over a simple sigmoid cline is slight for P3D05, the other SNP having a steep cline (AIC = 711.4 for stepped vs 712.3 for sigmoid). Of the SNPs with shallower clines, a stepped cline was strongly supported for P7G05 and P9D09 (809.4 vs 815.4, and 816.7 vs 833.3, respectively) but not for P7C08 (AIC=761.6 for stepped vs 762.0 for sigmoid).

### Estimating dispersal and selection

The four genetic loci with steep clines (*Eda*, *ATP1a1*, P3D05, and P6A10) are more or less fixed on either side of the zone and can be treated as diagnostic. We used these four loci to calculate a Hybrid Index (HI), as described in Barton & Gale (1993; hereafter BG93). The distribution of the HI through the hybrid zone is shown in Figure 5, while the mean and variance for each sampling location are given in Table 1.

**Figure 5.**
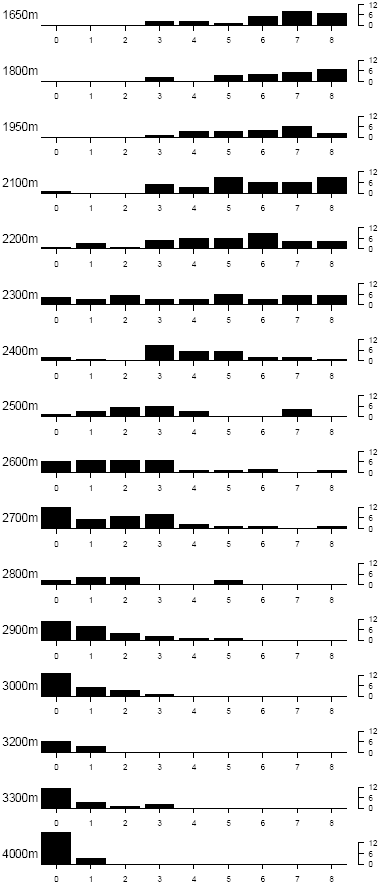
Bar plots of Hybrid Index through the hybrid zone. The Hybrid Index is given by the total number of anadromous alleles an individual carries at *Eda, ATP1a1*, P3D05, and P6A10.

The variance in HI in the center of the zone was used to estimate mean linkage disequilibrium 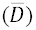; this parameter can then be used to estimate the dispersal rate. A portion of the variance in HI in the zone center arises from the variance in allele frequency at the individual loci (var(*p*) in BG93), and the remainder is due to migration bringing in purer genotypes. Taking the 64 individuals sampled at 2300m and 2400m (between which the clines at these four loci are roughly centered), we estimate the variance in allele frequency to be var(*p*) = 0.0025. From the first term of equation 2b of BG93, the proportion of 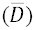 due to var(*p*) is 0.0309. The mean HI in this sample is 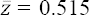, and the variance in HI is var(*z*) = 0.087. The remaining variance (0.0875-0.0309 = 0.0566) leads to the calculation 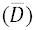 = 0.151, with 95% confidence intervals (using F_67,Infinity_) of 0.112 to 0.228.

Next, we convert the jointly fitted slope of these four clines (-2.86) to cline width via 4/abs(-2.86) = 1.39km (see Gay et al. 2008). We can use the estimates of 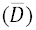 and cline width to estimate effective dispersal within the hybrid zone, as measured by the standard deviation *σ* of the distance between where a fish is hatched and where it reproduces, using the equation 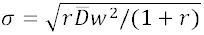 (p24 of BG93). Our samples were restricted to fish over 40mm, which were very likely sampled after they had dispersed. None of these four loci are physically linked, so *r* = 0.5, and thus *σ* = 0.25 km gen^-1/2^.

Finally, we can use dispersal and cline width to estimate *s*\*, the difference in mean fitness at that locus or trait between populations at the center of the zone and those at the edge, using *s*\* = (1.732*σ*/*w*)^2^ (p16 of BG93). We estimated *s*\* = 0.097 for the joint cline at *Eda*, *ATP1a1*, dorsal fin, 2^nd^ dorsal spine, left pelvic spine and peduncle depth.

## Discussion

In this paper we looked for evidence that strong direct selection could uncouple clines for loci and traits assumed to be under divergent ecological selection, thereby reducing the linkage disequilibria between them and impeding progress towards speciation. The clines for *Eda*, *ATP1a1*, dorsal fin length, caudal peduncle depth, pelvic spine length, and second dorsal spine length were found to be concordant and coincident. The cline for pectoral fin length was steeper and located 200m downstream from the other morphological traits. Interestingly, two anonymous SNPs (P3D05 and P6A10) also had steep clines located close to the joint cline for the selected traits and loci. The remaining eight SNPs had shallower clines with a wide range of slopes and centers (Table 2).

### Why do some clines have the same slope and centers?

The concordance and coincidence of the clines at *Eda*, *ATP1a1*, dorsal fin length, second dorsal spine length, pelvic spine length, and caudal peduncle depth suggests that these traits and loci are experiencing roughly the same selection regime, with similar intensities of direct selection. It also suggests that the position along the cline at which selection switches from favoring the marine allele to favoring the freshwater allele is also very similar. However, it is difficult to explain why they would all experience similar direct selection, since their functions are diverse and include body armor, osmoregulation and swimming capacity. An alternative explanation for the equivalent clines among such diverse traits and genes is that indirect selection is strong and similar on all of them because of strong linkage disequilibria (LD) between the loci and traits that pushes the clines to have the same slopes and centers (Barton 1983, Szymura & Barton 1991).

We can get an approximate estimate of the relative strengths of direct and total (i.e. direct + indirect) selection acting on a locus by fitting a stepped cline. In the center of the zone, LD between selected loci generates indirect selection on all of them together, steepening each of the clines (Szymura & Barton 1991). At the same time, an allele that has introgressed from the population on one side of the hybrid zone to the other side of the zone is at low frequency in the tail of the cline and is thus unlikely to be in LD there with introgressing alleles at other loci. The rate at which a focal allele moves further into the other population depends mostly on the relationship between dispersal and selection acting directly on that locus (and linked regions of the surrounding chromosome; Szymura & Barton 1991, Barton & Gale 1993, Baird 1995). Loci that are experiencing strong total selection but weak direct selection should therefore have a stepped cline, with a steep section in the center of the zone (where there is more LD) and shallower clines on either side.

The model fit for a stepped cline at *Eda* and *ATP1a1* was worse than the fit for a simple sigmoid cline. Similarly, a stepped cline was a worse fit for the four morphological traits in the joint cline (dorsal fin length, caudal peduncle depth, pelvic spine length, and second dorsal spine length). This is a puzzling finding. These clines are all steep compared to the estimated rate of dispersal, and highly concordant and coincident (Figures 2 and 3). There certainly is plenty of LD at the cline center: at 2300m, the average *D* between *Eda*, *ATP1a1*, and P3D05 is 0.10, and *D’* is 0.46, indicating that LD is 46% of its maximum possible value in this site (estimated with the R package ‘genetics’: R Core Team, 2014, Gregory Warnes et al., unpublished). Indirect selection is therefore expected to be acting in the center of the hybrid zone and not at the edges, and these six clines should be stepped.

**Figure. 2.**
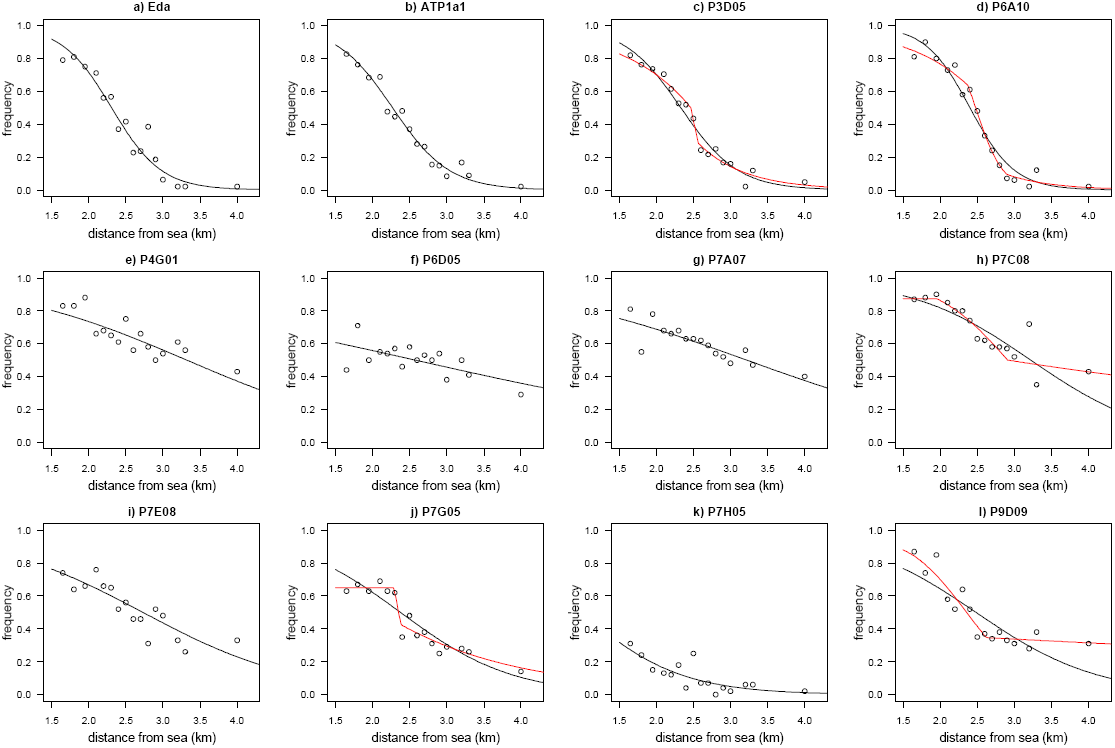
Individually fitted lines for the 12 genetic loci. The sigmoid cline is shown in black; the stepped cline is shown in red for the ci where this was a better fit.

**Figure 3a-e.**
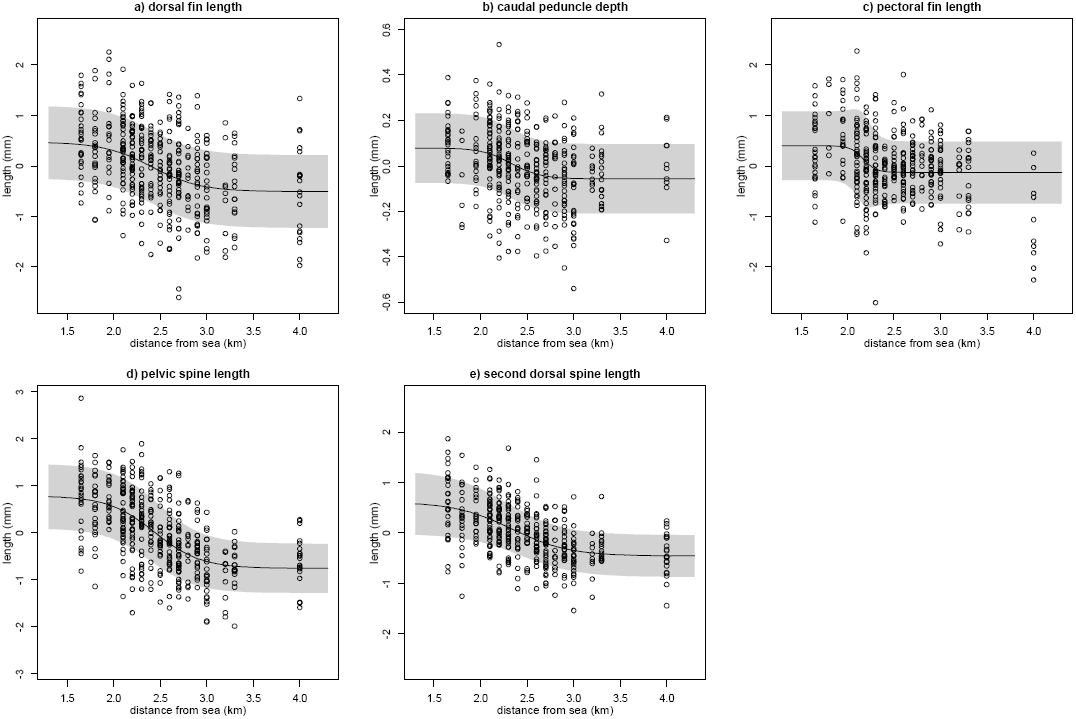
Clines in morphological traits through the hybrid zone. The width of the shaded band shows the trait variance through the zone, as calculated from *σ*_1_, *σ*_2_, and *σ*_3_.

One alternative explanation is that our assumption above is wrong: despite the dissimilarities in function, these loci and traits are actually experiencing similar amounts of direct selection (*s*\*∼0.1, see last section of Results), and their selection regimes all transition between favoring anadromous and freshwater alleles in the same part of the river (close to the estuary-freshwater ecotone). Linkage disequilibria between the loci generate indirect selection, but perhaps this additional indirect selection is only strong enough to bring the clines into full concordance and coincidence, and not strong enough to generate stepped clines. Modeling work on the behavior of strongly selected loci in hybrid zones would be a useful way to examine this scenario.

### Why do some clines have different centers and slopes?

We found that two anonymous SNPs had steep clines. The cline at P3D05 was very similar to the joint cline described above (centered at 2330m with a slope -2.59. The cline for P6A10 was slightly steeper (-3.3) and displaced about 100m upstream (at 2402m) from the joint cline, which is comparable to the dispersal distance (125 m). P3D05 is located at 10349kb along LGIII and is embedded within a gene called “amyloid beta (A4) precursor protein-binding, family B, member 1 interacting protein”. P6A10 is at 12601kb on LGVII in a gene called “mitochondrial calcium uptake 2”. Neither gene has been explicitly connected to freshwater adaptation in sticklebacks. Jones et al. (2012) did not find significant anadromous-freshwater divergence in the vicinity of either SNP in their genome scan. Hohenlohe et al. (2010) saw no significant divergence in that part of LGIII, although they did highlight five divergent SNPs (at 12212041, 12505622, 12525947, 12801733, and 12987907) in the vicinity of P6A10 on LGVII (Figure 8a in Hohenlohe et al. 2010, and P. Hohenlohe, pers. comm).

Interestingly, there is support for a stepped cline at P6A10 (Table 2a and Figure 2d). The most plausible explanation is that it is experiencing both some direct selection (*s*_*e*_ = tslope x *s\** = 0.11, see Szymura & Barton 1991) on either side of the hybrid zone and a combination of direct and indirect selection in the center (*s\** = 0.34). The source of this indirect selection must be linkage disequilibria with other selected loci, either on the same chromosome or elsewhere in the genome.

### The non-concordant cline for pectoral fin length

The cline for pectoral fin length is unlike any of the other morphological traits or genetic loci. The cline is centered between 2100 and 2200 m, a few hundred meters downstream of the other clines, and it is much steeper (-14.5 compared with c. -3 for the other morphological traits, Table 2b), with the caveat that the complicated distribution of phenotypes makes estimating cline shape quite difficult (Figure 3c and Figure S1). The cline position seems to arise because very few fish with relatively longer fins were sampled further upstream than 2100m. There is a deep (3m) pool at 2100m and the streambed salinity here is 20ppt at all points of the tidal cycle. So, even though it is clearly possible for fish with long pectoral fins to swim into the upper parts of the estuary, and many fish with otherwise anadromous morphology are found upstream of 2100m (Figure 3a, c-e), it appears that long finned fish restrict themselves to the lower, more brackish, parts of the creek.

Why might pectoral fin length show a different cline center than other traits? It is possible that shorter pectoral fins are highly deleterious in migrating anadromous fish, such that returning anadromous sticklebacks are all long-finned. If anadromous fish only migrate into the lower part of the estuary then the cline for pectoral fin length will coincide with that limit. This scenario implies that there is a link between the phenotype of a fish and its preferred location in the river (see below). Fish with otherwise anadromous morphology found further upstream might remain in the estuary year round, or overwinter in the sea close to the mouth of the creek. While we do not currently know the migratory limit in the estuary for the anadromous population, it would be possible to use stable isotope ratios to distinguish migratory from non-migratory fish (e.g. Arai et al. 2003).

An additional explanation for the downstream position of the pectoral fin cline is that the stream alleles for the underlying loci are dominant, such that a genetically intermediate hybrid would have the short pectoral fin phenotype. However, lab crossed F1 hybrids between stream and anadromous fish from Bonsall Creek showed no net dominance for pectoral fin area (Figure 3B in Dalziel et al 2012).

### The structure of the Bonsall Creek hybrid zone

One puzzling feature of this hybrid zone is that the observed clines are generally very steep compared to the movement capability of the fish. A mark-recapture study conducted on 23^rd^ and 24^th^ May 2006 (T. Vines & A. Dalziel, unpublished data) found several fish that had moved several hundred meters overnight. If fish move tens of meters per day in a random direction, selection would need to be extremely strong to maintain these steep clines: almost all fish crossing the hybrid zone would need to be eliminated before they could be sampled. It is difficult to imagine any selective force that could accomplish this. Adult sticklebacks are able to cope with a wide range of salinities without suffering large effects (e.g. Schaarschmidt et al. 1999, Barrett et al. 2009), so the salinity differences between the estuary and freshwater cannot be a big enough source of mortality. Moreover, piscivorous birds and fish are present throughout the hybrid zone, it seems very unlikely that their selective impacts would be restricted to just one side of the zone. The density of predators would also need to be very high to take out the constant flux of immigrants.

A more reasonable explanation is that while fish could easily traverse the zone, they instead choose a point along the river according to their genotype and phenotype and stay there over the breeding season. This must be the case for the anadromous fish, which spend the winter in the sea and migrate into the river in the spring (Hagen 1967). The Bonsall Creek hybrid zone is thus similar to hybrid zones between migratory and resident bird species (e.g. Rohwer et al. 2001, Brelsford & Irwin 2009) where one species migrates long distances over the winter and the zone is only reformed in the breeding season. The key parameter in these zones is not the total distance travelled (which is orders of magnitude greater than the zone width), and instead is the distance between where the organism is born and where it reproduces. The standard deviation in the latter distance is estimated by *σ* = 0.25 km gen^1/2^. above: in this hybrid zone, an individual reproduces an average of 125m upstream or downstream of where it was born.

The steep concordant clines in Bonsall Creek perhaps arise because breeding location is determined by a continuous mating habitat preference, and the preference loci are in tight linkage disequilibria with the traits and loci we examined here. In this scenario, pure anadromous fish prefer to nest in the saltiest part of the river near the sea, and fish with an increasing proportion of stream alleles at the preference loci are found closer and closer to freshwater. The strong LD between the alleles underlying habitat preference and the stream or anadromous alleles for other morphological or genetic clines should be maintained by strong total selection. Alternatively, there may be no segregating variance for habitat preference, and the fish may follow a simple rule like ‘remain in the part of the river where the effort to maintain osmotic balance and swimming effort is minimized’. What is the source of this selection? Unlike adults, eggs and newly hatched offspring cannot change their location within the river, and must therefore experience the changing salinities associated with the tidal cycle. The effects of salinity on egg hatching success and later fish growth in stickleback appear to be significant. For example, the hatching success of eggs from Belgian freshwater populations is better in low than high salinity, while the converse is true for marine populations (Heuts 1947). More recent studies have found low hatching success and growth rate of freshwater fish in higher salinities (Marchinko & Schluter, 2007; DeFaveri and Merila 2014).

Interestingly, some hybrid zones between anadromous and stream sticklebacks are found exclusively in fresh water, such as on tributaries of the nearby Fraser River (Virgil & McPhail 1994, Dalziel et al. 2012). Since all of these hybrid zones are similarly narrow, it seems unlikely that adaptation to different salinities in the early life stages is the major source of selection against hybrids or against the offspring of fish breeding in the ‘wrong’ part of the river. Instead, as hypothesized by Hagen (1967), it seems plausible that the majority of selection occurs over the winter, and is largely against hybrids that are unfit in both the stream or the sea. Because linkage disequilibria are generally quite strong in this hybrid zone, hybrids will tend to have many features that make them poorly suited to surviving in either environment. For example, a hybrid that remains in the stream over the winter will need to cope with the combination of low temperatures and fresh water (which in combination can be challenging for anadromous fish, see Heuts 1947, Schaarschmidt et al. 1999), and the need to efficiently maintain position in the stream. It will also need to compete effectively with pure stream sticklebacks for (presumably) limited food resources. By contrast, hybrid fish in the marine environment will need to cope with the higher density of vertebrate predators, and compete for food with the pure anadromous fish. Studies with sympatric stickleback species pairs have shown that selection against hybrids via competition for food can be strong (Rundle 2002), particularly when coupled with predation (Vamosi & Schluter 2002, Rundle et al. 2003). It is also possible that some hybrids do not choose between the stream or marine environment at all, and overwinter in the more intermediate environment of the estuary.

## Conclusions

This study looked for evidence that direct selection could uncouple clines at loci known to be under ecological selection. We examined two well-studied genes in a hybrid zone between stream and anadromous sticklebacks, and found that the clines at *Eda* and *ATP1a1* shared the same slope and center, as did two other anonymous SNPs and four morphological traits. The concordance and coincidence of these clines suggests that indirect selection is the major force in determining their shape and location, and that direct selection on the individual genes and traits has not been sufficient to uncouple them. The cline for pectoral fin length was much steeper and centered 200m downstream, perhaps because long-finned anadromous fish do not migrate any further into the creek. Given the importance of pectoral fins for steady locomotion in sticklebacks, pectoral fin length may be under strong selection in migrating anadromous fish.

## Acknowledgments

The authors are grateful to Taylor Gibbons for help in the lab, Jeremy Schmutz for information on the SNP loci, Jessica Hill and Chrissy Spencer for help with sample collection, and to James Thomas and other members of the Halalt first nation for access to their land and information on the history of the creek. We thank Nick Barton, Sean Rogers and Thomas Lenormand for helpful discussions, while Rowan Barrett and Dave Toews provided comments on an earlier version. This work was supported by the Natural Sciences and Engineering Research Council of Canada (NSERC) through Discovery and Discovery Accelerator grants to PMS, Discovery and Special Research Opportunity grants to DS, and a Canada Graduate Scholarship to ACD and AA. THV was supported by a European Union FP6 Outgoing International Fellowship and a real job. TV was supported by the NSERC-CREATE Training Program in Biodiversity Research and NSF grant IOS-1145468.

